# Self-organization of vascularized skeletal muscle from bovine embryonic stem cells

**DOI:** 10.1101/2024.03.22.586252

**Authors:** Marina Sanaki-Matsumiya, Casandra Villava, Luca Rappez, Kristina Haase, Jun Wu, Miki Ebisuya

## Abstract

Cultured beef holds promising potential as an alternative to traditional meat options. While adult stem cells are commonly used as the cell source for cultured beef, their proliferation and differentiation capacities are limited. To produce cultured beef steaks, current manufacturing plans often require the separate preparation of multiple cell types and intricate engineering for assembling them into structured tissues. In this study, we propose and report the co-induction of skeletal muscle, neuronal, and endothelial cells from bovine embryonic stem cells (ESCs) and the self-organization of tissue structures in 2- and 3-dimensional cultures. Bovine myocytes were induced in a stepwise manner through the induction of presomitic mesoderm (PSM) from bovine ESCs. Muscle fibers with sarcomeres appeared within 15 days, displaying calcium oscillations responsive to inputs from co-induced bovine spinal neurons. Bovine endothelial cells were also co-induced via PSM, forming uniform vessel networks inside tissues. Our serum-free, rapid co-induction protocols represent a milestone toward self-organizing beef steaks with integrated vasculature and innervation.

## Introduction

Cultured meat, also referred to as cultivated meat, cell-based meat, lab-grown meat, or clean meat, is produced through culturing animal skeletal muscle cells in vitro^1–4^. Cultured meat has the potential to address ethical, environmental, and health concerns associated with conventional meat production from livestock.

Current practices in cultured beef production predominantly rely on myosatellite cells and mesenchymal stem cells (MSCs) isolated from adult bovine tissue biopsy and bone marrow samples^4–6^. The proliferation capacity of these adult stem cells diminishes over long-term culture. Although they can be immortalized^7^, the genetic modifications required for immortalization may present a challenge in obtaining food authorization. The differentiation capacity of adult stem cells is also limited. For instance, myosatellite cells are committed to muscle fate; although MSCs can form osteoblasts, adipocytes, and chondrocytes in addition to myocytes, the differentiation trajectory of MSCs is still restricted and dependent on the tissue source from which they are isolated. By contrast, pluripotent stem cells, namely embryonic stem cells (ESCs) and induced pluripotent stem cells (iPSCs) proliferate indefinitely, offering a more stable cell source for cultured meat. Pluripotent stem cells can also differentiate into cells of all three germ layers: mesoderm, ectoderm, and endoderm. Unmodified ESCs are more favorable for food production than iPSCs that need genetic reprogramming. The derivation of stable bovine ESCs was historically challenging but recently achieved^8^, and a large-scale culture of bovine ESCs in stirred bioreactors has been reported^9^.

Another challenge in the cultured meat field is how to produce structured steaks instead of ground meat. The incorporation of diverse cell types within the muscle tissue holds the key to more accurate recreation of steaks^10,11^. For in vitro tissues to keep growing, oxygen and nutrient supply through blood vessels or alternative porous structures is also crucial^12^. The current manufacturing strategy for cultured beef steaks involves the separate preparation of distinct cell types followed by their assembly. This assembly process necessitates advanced engineering techniques, such as bioprinting purified cells with a bioink, co-culturing cells on edible porous scaffolds, and stacking myocyte-laden hydrogel modules^13–16^. Additionally, electrical or mechanical stimuli are often applied to facilitate muscle maturation and improve the texture^16–21^. While these engineering approaches are promising and evolving rapidly, they add extra manufacturing steps to cultured steak production. A potential alternative approach is utilizing pluripotent stem cells and leveraging their differentiation and self-organization abilities that have been demonstrated by the development of a variety of complex organoids in recent years^22–28^.

In this study, we utilized bovine ESCs to co-induce skeletal muscle, neuronal, and endothelial cells, fostering the self-organization of muscle tissues with innervation and vascularization.

## Results

### Induction of PSM and muscle fibers from bovine ESCs

We first established a protocol to induce muscle fibers from bovine ESCs (Fig 1a; Supplementary Fig. 1). As skeletal muscle is generated from presomitic mesoderm (PSM) through transient structures of somites and dermomyotomes in embryonic development^29^, our stepwise protocol comprised the induction of PSM followed by the induction of myoblasts and myocytes. Based on the PSM induction protocol we previously reported^30–33^, bovine ESCs were treated with CHIR (WNT signaling activator), bFGF, SB431542 (TGFb signaling inhibitor), and DMH1 (BMP signaling inhibitor) for 2 days. The expression levels of pluripotency markers, *NANOG*, *OCT4*, and *SOX2*, decreased before day 2 (Supplementary Fig. 2a) while those of a mesodermal marker *BRACHYURY* (*BRA*, also known as *T*) and a PSM marker *TBX6* increased (Supplementary Fig. 2b). Most cells were TBX6-positive on day 2, suggesting efficient PSM induction (Fig. 1b). Since the remarkable characteristic of PSM cells is the segmentation clock, the oscillatory gene expressions that regulate the timing of somite formation, we monitored the expression pattern of HES7, a core gene of the molecular oscillator^34,35^. A HES7 promoter-luciferase reporter^30–33^ showed an elevated expression around day 2 (Fig. 1c, top), simultaneously displaying clear oscillations of the segmentation clock (Fig 1c, bottom). Consistent with the previous report^30^, the oscillation period of the bovine segmentation clock was ∼4 hours. These results confirmed the efficient induction of bovine PSM from bovine ESCs.

**Figure 1.**
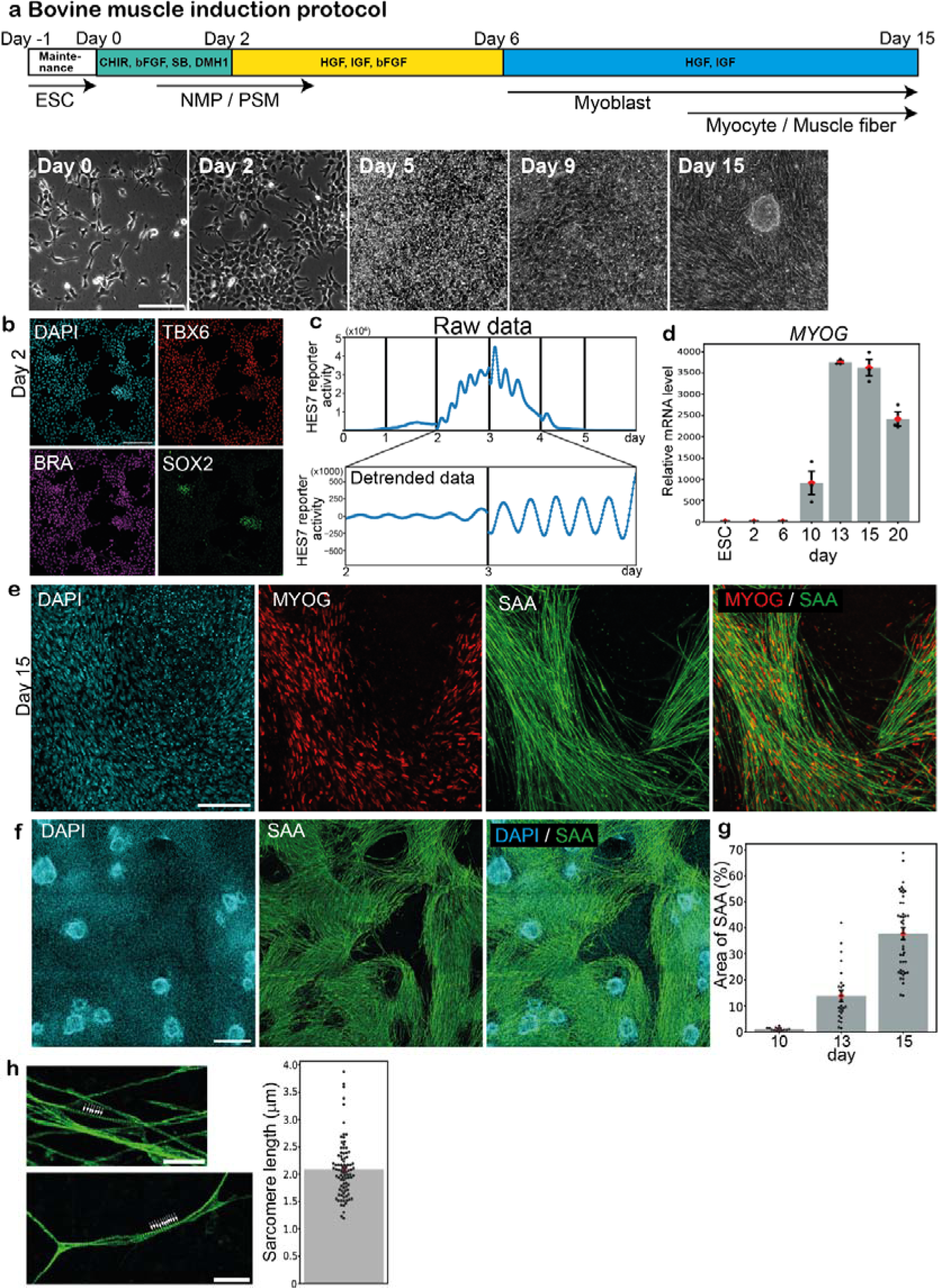
Induction of PSM and muscle fibers from bovine ESCs. **a,** Induction protocol for bovine myocytes and muscle fibers from bovine ESCs. PSM cells were initially induced from ESCs using CHIR, bFGF, SB, and DMH1. Then myocytes were induced from PSM using HGF, IGF, and bFGF. Maintenance: Maintenance medium, ESC: Embryonic stem cell, NMP: Neuromesodermal progenitor, PSM: Presomitic mesoderm. Typical bright field images are also shown (bottom). **b,** IHC images of the day-2 culture. TBX6 is a PSM marker. SOX2- and BRA-double positive cells are NMPs. 3 independent experiments showed similar staining patterns. **c,** The oscillatory activity of the HES7 promoter-luciferase reporter in PSM. Raw (top) and detrended (bottom) signals. The vertical lines indicate the timings of medium change, which resets the oscillation phase. 3 independent experiments showed similar oscillation patterns. **d,** qRT-PCR quantification of *MYOG* expression. The average value of ESC sample was set to 1. Mean ± SEM. N = 3 from 3 independent experiments. **e,f,** IHC images of the day-15 cultures. MYOG and SAA are myocyte/muscle fiber markers. 4 independent experiments showed similar staining patterns. **g,** Percentage of the SAA-positive area in the field of view. Mean ± SEM. N = 14 (day 10), 25 (day 13), and 42 (day 15) from 3-9 independent experiments. **h,** Sarcomeric structures on SAA-positive muscle fibers (arrowheads in the left pictures) on day 15. The sarcomere length was defined as the distance between intensity peaks (right). Mean ± SEM. N = 88 from 4 independent experiments. Scale bars: 150 µm (a,b,e), 300 µm (f), and 30 µm (h). Microscopes: IX73 (a) and FV3000 confocal (b,e,f,h).

To further induce skeletal muscle from bovine PSM, we adapted the muscle induction protocol previously reported for mouse and human cells^36–38^. Bovine PSM cells were treated with HGF, IGF, and bFGF for 4 days before further treatment with HGF and IGF for 9 days (Fig. 1a). The expression levels of myoblast markers, *MYF5* and *MYOD1*, increased around day 6 (Supplementary Fig. 2c). The expressions of myocyte markers, *MYOG, MYF6*, *MYH7*, and *MYH4*, were induced around day 10 (Fig. 1d; Supplementary Fig. 2d). The MYOG- and sarcomeric α-actinin (SAA)-positive muscle fibers first appeared around day 10 and peaked around day 14 (Fig. 1e-g). These bovine muscle fibers displayed sarcomeric structures, and their average sarcomere length was 2.1 µm (Fig. 1h), similar to previously reported values for in vivo bovine skeletal muscles^39,40^. The ∼15-day duration of bovine myocytes and muscle fiber induction was slightly shorter than the one described in the human skeletal muscle induction protocol^37^, potentially reflecting the faster pace of bovine embryonic development compared with human^30^. These results collectively established the induction protocol for bovine myocytes and muscle fibers from bovine PSM. Note that the protocol does not require serum.

### Co-induction of bovine muscle and spinal neurons

The induced bovine myocytes and muscle fibers displayed spontaneous fluctuations of intracellular calcium levels (Fig. 2a,b; Supplementary Video 1). The calcium pulses were effectively suppressed by the treatment with curare (Fig. 2b,c; Supplementary Video 1), an acetylcholine receptor blocker that is used to inhibit neuromuscular junctions^28,41,42^. This suggested the presence of neuromuscular junctions connecting skeletal muscles and spinal neurons. Indeed, we detected TUJ1-positive neurons intermingling with SAA-positive muscle fibers in the day-15 muscle culture (Fig. 2d,e). A neurofilament maker SMI-32 and another muscle marker, myosin heavy chain (MHC), showed similar staining patterns of aligned neurons and muscle fibers (Fig. 2f,g). The expression of a neuronal marker *SOX2* also increased around day 10-13 (Supplementary Fig. 2a). These neurons could be derived from neuromesodermal progenitors (NMPs), given that the day-2 PSM culture contained a fraction of SOX2- and BRA-double positive NMPs (Fig. 1b; Supplementary Fig. 1b). These results suggested that spinal neurons were co-induced from bovine ESCs, transmitting a signal to the skeletal muscles through neuromuscular junctions. The stimulation from neurons may be beneficial for muscle maturation, potentially offering an alternative to the external electrical and mechanical stimuli currently employed in the muscle bioengineering field^16–21^.

**Figure 2.**
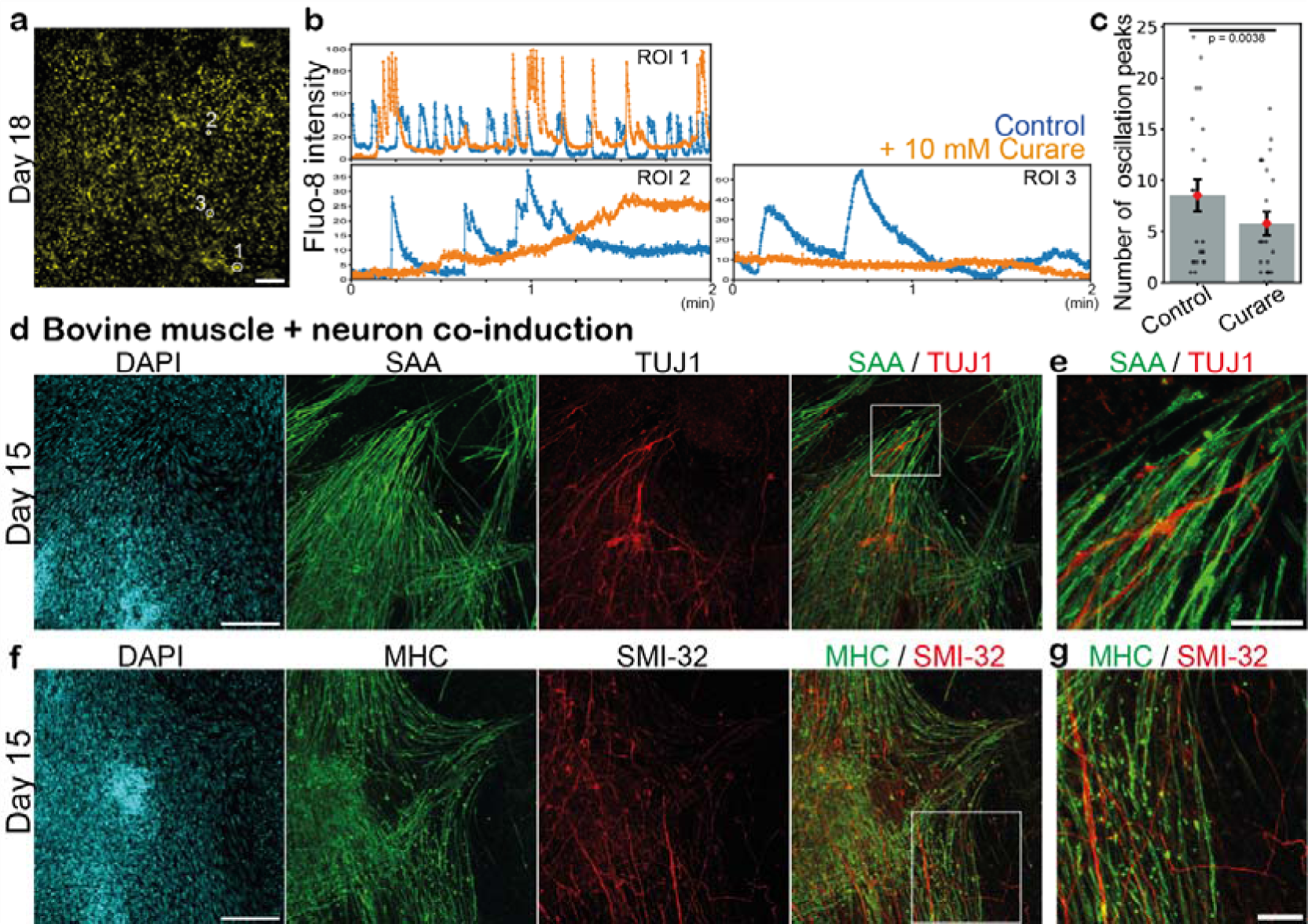
Co-induction of bovine muscle and spinal neurons. **a,** Fluo-8 calcium signal of the day-18 culture. **b,** Calcium level fluctuations in the absence (blue) and presence (orange) of 10 µM curare, an acetylcholine receptor blocker. Regions of interest (ROIs) are marked in a. **c,** Quantification of the total number of calcium oscillation peaks within 2 min in the absence and presence of curare. Mean ± SEM. N = 23 (control) and 23 (curare) from 3 independent experiments. P-value is from paired t-test. **d-g,** IHC images of the day-15 cultures. TUJ1 and SMI-32 are neuronal markers. SAA and MHC are skeletal muscle markers. 3-8 independent experiments showed similar staining patterns. **e,g,** Enlarged images of d and f, respectively. Scale bars: 150 µm (a,d,f) and 50 µm (e,g). Microscopes: Thunder (a) and FV3000 confocal (d-g).

### Co-induction of bovine muscle and endothelial cells

We next developed a protocol to co-induce skeletal muscle and endothelial cells from bovine ESCs (Fig. 3a). The lack of blood vessels is a major challenge that currently hampers the development of large, mature organoids in the stem cell biology field. As blood vessels are first formed by endothelial cells, we prioritized the induction of bovine endothelial cells. Note that while endothelial cells arise primarily from lateral plate mesoderm in embryonic development^43,44^, a contingent of endothelial cells are derived from PSM and somites^44–49^. We thus hypothesized that endothelial cells and myocytes could be co-induced from the same PSM cell population. The BMP inhibitor DMH1 that prevents lateral plate mesoderm differentiation was kept in the protocol (Fig. 3a). On day 2, we added the angiogenic factors VEGF and forskolin (cAMP signaling activator) to the bovine PSM culture and initiated the induction of endothelial cells, adapting the reported protocols for human endothelial cells^50–52^. The elongated cells, positive for an endothelial marker VE-cadherin (VE-cad, also known as CDH5), appeared around day 8 and peaked around day 14 (Fig. 3b). The induced bovine endothelial cells formed interconnected vessel-like networks: the numbers of vessel segments and branching points increased during days 8-15 (Fig. 3c). The endothelial networks and muscle fibers tended to form small individual domains (Fig. 3d,e). The skeletal muscle domains encased by endothelial cells measured 200-300 µm in diameter on day 15 (Fig. 3f), which surpassed the intercapillary distance of approximately 100 µm in tissues^53^, yet remained within a comparable range. These results demonstrated the co-induction of skeletal muscle and endothelial cells from bovine PSM and the self-organization of vascular networks in 2-dimensional (2D) cultures.

**Figure 3.**
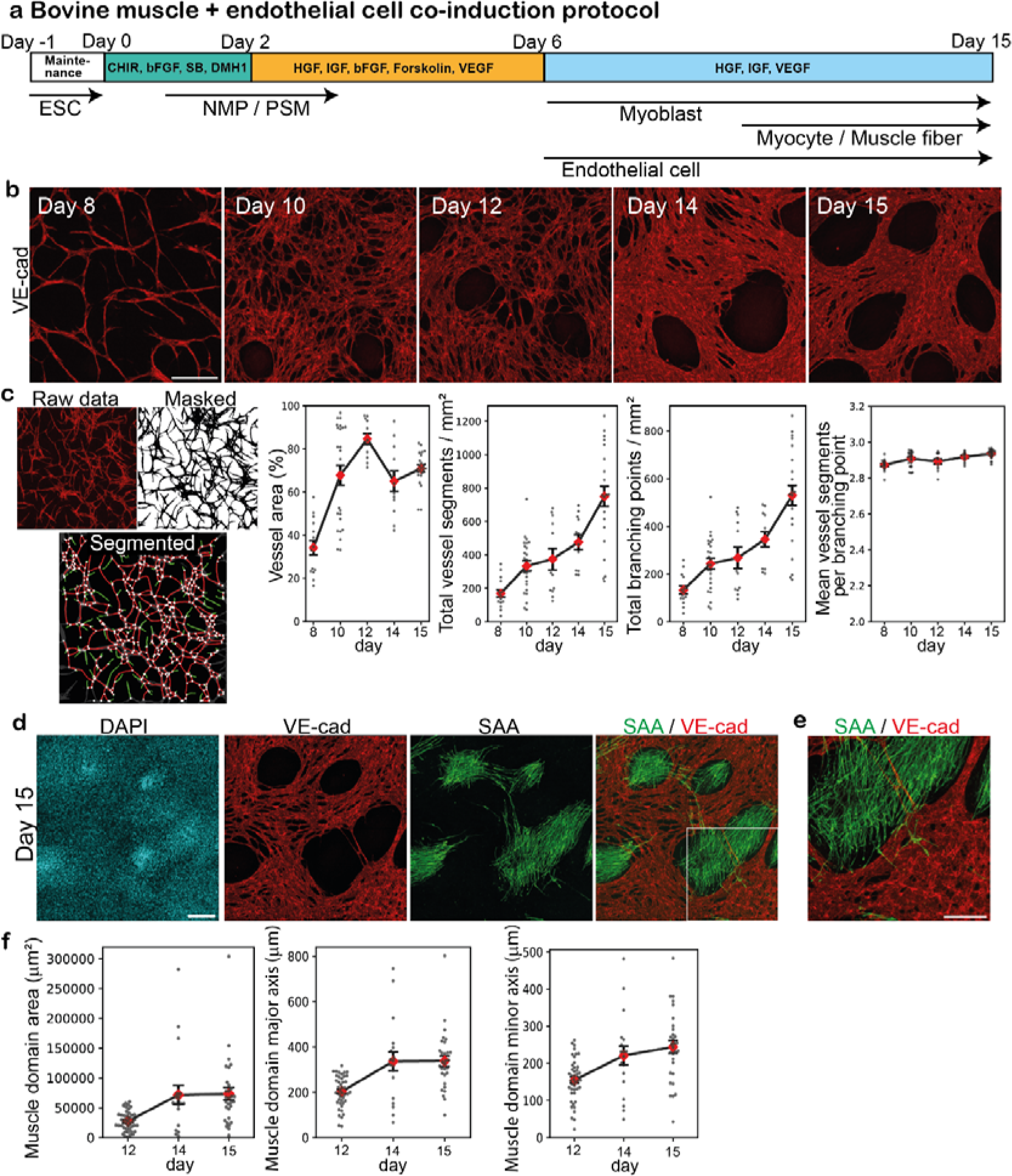
Co-induction of bovine muscle and endothelial cells. **a,** Co-induction protocol for bovine muscle fibers and endothelial cells. Forskolin and VEGF were added to induce endothelial cells from bovine PSM cells. **b,** IHC images over the time course. VE-cad is an endothelial cell marker. 3-6 independent experiments showed similar staining patterns. **c,** Characterization of the endothelial cell networks using VascuMap^68^. After the IHC images were masked and segmented, the vessel segments (red edges) and branching points (white nodes) were identified and quantified using the connectivity analysis pipeline. The vessel segments that do not connect two points (green edges) were not counted. The vessel area, the total number of vessel segments, the total number of branching points, and the mean number of vessel segments per branching point were quantified. Mean ± SEM. N = 15 (day 8), 25 (day 10), 12 (day 12), 12 (day 14), and 23 (day 15) from 3-6 independent experiments. **d,** IHC images of the day-15 culture showing endothelial and skeletal muscle domains. 6 independent experiments showed similar staining patterns. **e,** Enlarged image of d. **f,** Size of skeletal muscle domains. The area, major axis length, and minor axis length of VE-cad-negative domains were quantified. Mean ± SEM. N = 43 (day 12), 19 (day 14), and 30 (day 15) from 3-4 independent experiments. Scale bars: 300 µm (b,d) and 150 µm (e). Microscope: FV3000 confocal.

### Induction of bovine muscle fibers in 3D cultures

With the ultimate goal of self-organizing beef steaks in mind, we applied our bovine ESC differentiation protocols to 3-dimensional (3D) cultures. We developed a protocol to induce bovine muscle fibers in 3D cell aggregates (Fig. 4a). Bovine PSM cells were induced according to our 2D protocol, and on day 2, a 3D aggregate was created out of 20,000 PSM cells (Fig. 4b), which contained a small fraction of NMPs. The 2% Matrigel was added to the medium to bolster the structural integrity of aggregates: without Matrigel, the aggregates gradually fell apart, and a higher concentration of Matrigel triggered the separation of two aggregate domains (Supplementary Fig. 3). The PSM cell aggregates were treated with the skeletal muscle-inducing cocktail, and after day 6, they were cultured on a rotary shaker that facilitated the diffusion of oxygen and nutrients (Fig. 4a). The aggregates were spherical and ∼600 µm in diameter (Fig. 4c), and the size did not dramatically change over the time course, probably reflecting a declined cell proliferation during the muscle differentiation phase. SAA-positive muscle fibers were induced around day 10 (Fig. 4d; Supplementary Video 2). Consistent with the results from our 2D cultures, TUJ1-positive neurons were co-induced in aggregates, forming neuronal domains next to muscle domains (Fig. 4d,e; Supplementary Video 2). To assess the effects of aggregate size on cell fates, we prepared diverse sizes of aggregates by changing the initial PSM cell number. The proportions of the skeletal muscle and spinal neurons gradually decreased as the aggregate size increased (Fig. 4f,g). These results established the induction protocol for a bovine 3D tissue comprising skeletal muscle and spinal neurons from bovine ESCs.

**Figure 4.**
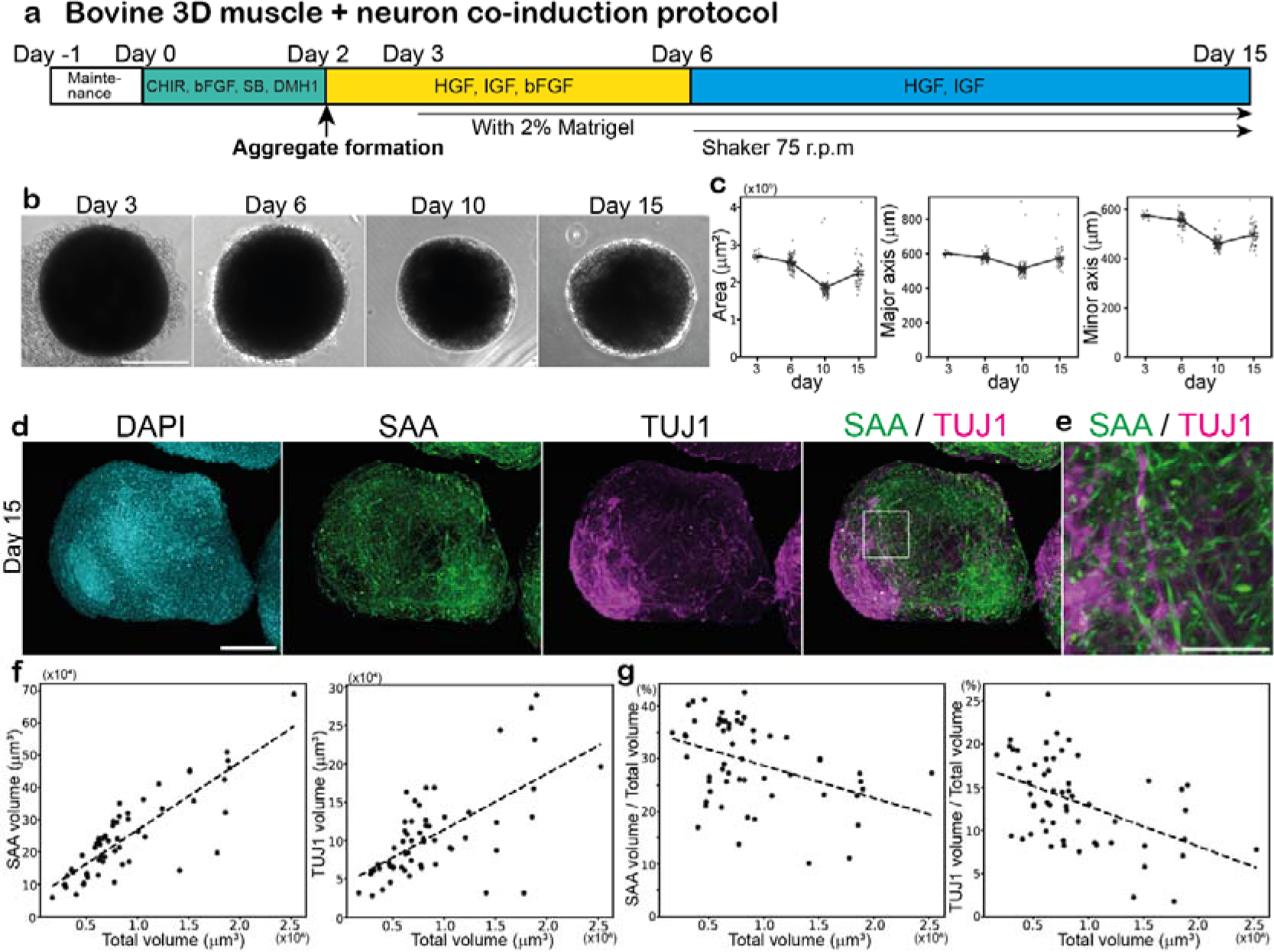
Induction of bovine muscle fibers in 3D cultures. **a,** 3D induction protocol for bovine muscle fibers and spinal neurons. The aggregate of 20,000 bovine PSM cells was made on day 2. Matrigel was added to the medium on day 3. Aggregates were cultured on a rotary shaker after day 6. **b,c,** Characterization of aggregates over the time course. The area, major axis length, and minor axis length of the aggregates were quantified using 2D bright field images. Mean ± SEM. N = 15 (day 3), 93 (day 6), 90 (day 10), and 48 (day 15) from 3 independent experiments. **d,** IHC images of the day-15 aggregate. SAA and TUJ1 are muscle fiber and neuronal markers, respectively. 6 independent experiments showed similar staining patterns. **e,** Enlarged image of d. **f,g,** Effects of aggregate size on cell fates. The aggregates were made from 10,000-30,000 PSM cells on day 2, and the percentages of the SAA- and TUJ1-positive volumes were quantified in the day-15 aggregates. N = 57 from 7 independent experiments. Scale bars: 300 µm (b), 150 µm (d) and 50 µm (e). Microscopes: IX73 (b) and Light-sheet (d,e).

### Co-induction of bovine muscle and endothelial networks in 3D

Finally, we developed a protocol to co-induce muscle fibers and endothelial cell networks from bovine ESCs in 3D cultures (Fig. 5a). The aggregates of bovine PSM cells were created on day 2, and then treated with the induction cocktail for skeletal muscle and endothelial cells. The aggregate size was again ∼600 µm in diameter over the entire time course (Fig. 5b,c). VE-cad-positive endothelial cells emerged around day 10, forming interconnected vessel-like networks adjacent to the domains of muscle fibers inside the aggregates (Fig. 5d-f; Supplementary Video 3). ZO-1 staining unveiled similarly connected networks, suggesting the presence of tight junctions in the endothelial networks (Fig. 5g,h; Supplementary Video 4). Note that the endothelial networks displayed a regular pattern throughout an aggregate, penetrating deeply into the tissue (Supplementary Video 3,4). The abundant and relatively uniform distribution of vasculature will be advantageous for facilitating oxygen and nutrient delivery throughout the muscle aggregates, even though vascular connectivity via intraluminal flow remains to be seen. The endothelial network formation depended on the induction protocols: when PSM cells were induced without the BMP inhibitor DMH1, the subsequent endothelial cells did not form uniform networks, and the shape of the aggregates was no longer spherical (Supplementary Fig. 4; Supplementary Video 5). Unlike the aggregates that lack endothelial cells (Fig. 4g), the proportions of the muscle domain and the endothelial networks slightly increased or remained constant as the aggregate size increased (Fig. 5i,j). These results underscored the feasibility of generating vascularized skeletal muscle tissues from bovine ESCs.

**Figure 5.**
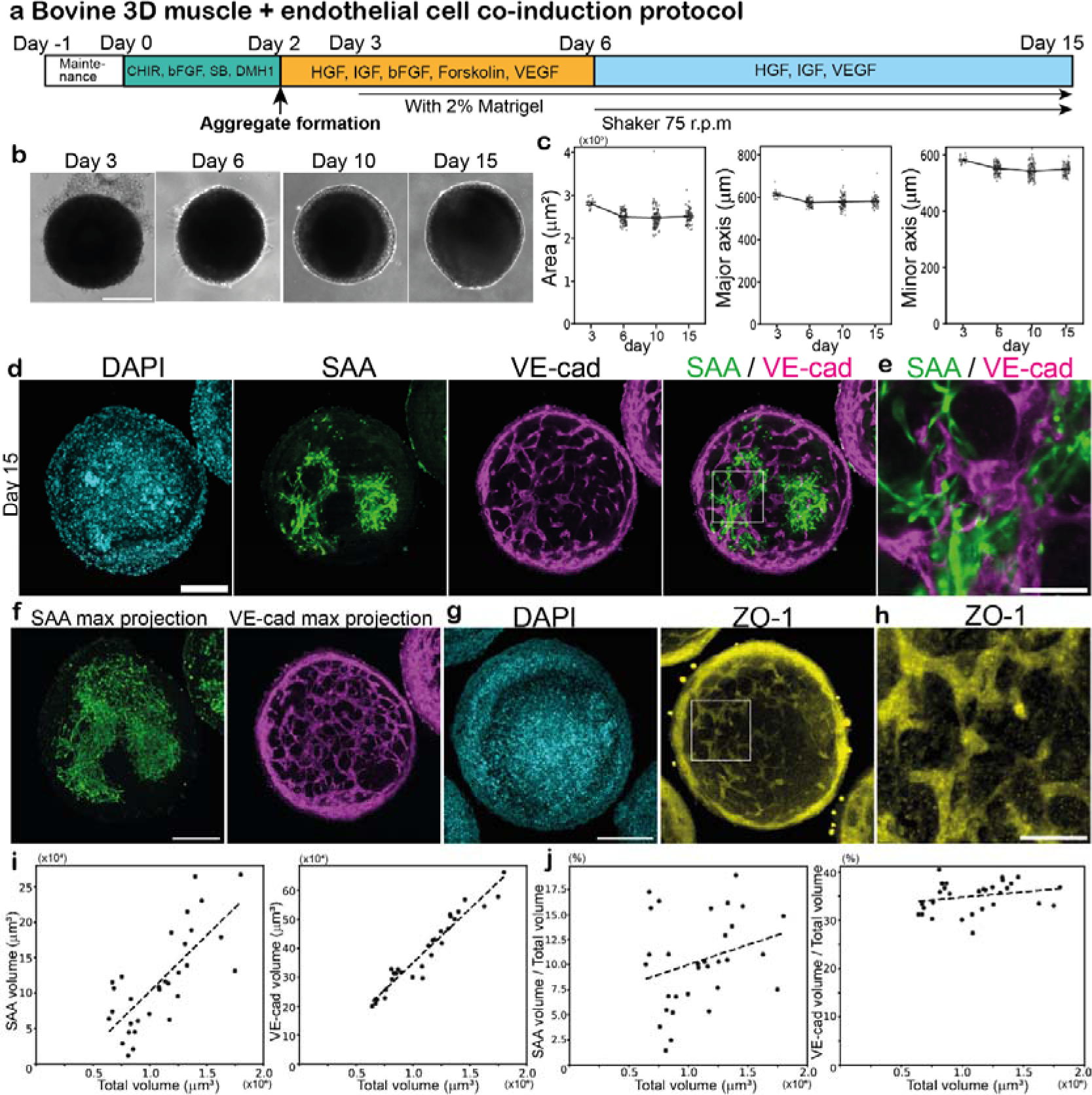
Co-induction of bovine muscle and endothelial networks in 3D. **a,** 3D co-induction protocol for bovine muscle fibers and endothelial cell networks. The aggregate of 20,000 bovine PSM cells was made on day 2. Forskolin and VEGF were added to induce endothelial cells from PSM cells. **b,c,** Characterization of aggregates over the time course. The area, major axis length, and minor axis length of the aggregates were quantified using 2D bright field images. Mean ± SEM. N = 15 (day 3), 94 (day 6), 93 (day 10), and 65 (day 15) from 3 independent experiments. **d-h,** IHC images of the day-15 aggregates. **d,** SAA and VE-cad are muscle fiber and endothelial cell markers, respectively. 10 independent experiments showed similar staining patterns. See also Supplementary Video 3. **e,** Enlarged image of d. **f,** Max projection images of d. The max projection of SAA used all Z-stacks. The max projection of VE-cad used the Z-stacks within the 40 μm range around the central slice. **g,** ZO-1 is a tight junction marker. Max projection used the Z-stacks within the 40 μm range around the central slice. 3 independent experiments showed similar staining patterns. See also Supplementary Video 4. **h,** Enlarged image of g. **i,j,** Effects of aggregate size on cell fates. The aggregates were made from 10,000-30,000 PSM cells on day 2, and the percentages of the VE-cad- and SAA-positive volumes were quantified in the day-15 aggregates. Mean ± SEM. N = 31 from 7 independent experiments. Scale bars: 300 µm (b), 150 µm (d,f,g), and 50 µm (e,h). Microscopes: IX73 (b) and Light-sheet (d-h).

## Discussion

In this study, we demonstrated the potential of utilizing bovine ESCs as a stable cell source for cultured beef and steaks. The induction of bovine myocytes and muscle fibers from indefinitely proliferative stem cells should make cultured beef production more scalable and robust. A large-scale culture of bovine ESCs in bioreactors has already been established^9^. The induction of distinct cell types, such as endothelial cells and neurons, from the same cell source can also simplify the production process. All the protocols described in this study are serum-free and can be completed within 15 days. The rapid induction should be advantageous for potential applications. By contrast, the high costs of the growth factors, chemical compounds, and Matrigel used in the protocols present a future challenge. Cost reduction through the mass production of these materials or the development of plant-based alternatives will be the key.

Moreover, we generated a bovine 3D tissue containing skeletal muscle, spinal neurons, and endothelial cell networks. Several 3D skeletal muscle organoids were previously developed from human pluripotent stem cells, but not from bovine stem cells^28,54–56^. Even though vascularized skeletal muscle tissues were previously engineered, the muscle cells and endothelial cells were separately prepared and assembled^13,56–58^, or the cells were induced through the overexpression of master regulators^59^. Thus, the co-induction of bovine myocytes and endothelial cells without genetic modification as well as the subsequent self-organization of muscle fibers and endothelial networks inside the tissue represent a crucial step toward the production of structured beef steaks. The stimulation of skeletal muscle by spinal neurons should also facilitate the maturation of the muscle tissue. However, further functionalization of the bovine blood vessels and neuromuscular junctions will be necessary. As the lumen formation in blood vessels is stimulated by intraluminal flow, the next crucial step should involve establishing the connection between the endothelial networks within the muscle aggregates and in vitro perfusable vasculatures^60–63^. Perfusion through blood vessels is vital for nurturing steak-sized tissues, and perfusion with blood substitutes may improve the flavor of cultured steaks. As the blood vessel formation is also heavily influenced by stroma cells, such as pericytes and fibroblasts, co-inducing bovine stroma cells together with endothelial cells may prove beneficial^26,64,65^. Similarly, for the maturation of the neuromuscular junctions, further optimization of the induction protocol and longer-term culture may be necessary^28^.

Despite the remaining challenges, we propose the co-induction of multiple cell types and self-organization of tissues as feasible directions in the production of cultured beef steaks. Our approach will also be complementary to engineering approaches: co-induced cells can be mixed with a bioink and readily used for bioprinting, while muscle aggregates featuring endothelial networks can serve as building blocks for complex tissue assembly. Diverse types of cultured beef will offer novel food options.

## Methods

### Bovine ESC culture

The bovine ESC line was obtained from Dr. Juan Carlos Izpisua Belmonte^8^. The HES7 promoter-luciferase reporter bovine ESC line was described previously^30^. The ESCs were maintained without feeder cells and cultured on Matrigel (Corning, 356231)-coated dishes. Either mTeSR1 medium (STEMCELL Technologies) or StemFit medium (Ajinomoto, StemFit Basic04CT) supplemented with 20 ng/mL bFGF (PeproTech, AF-100-18B), 20 ng/mL Activin A (R&D, 338-AC), and 2.5 μM IWR1 (Tocris, 3532) was used as the maintenance medium. The ESCs were passaged every other day. The cells were trypsinized into single cells by TrypLE solution (Gibco, A1285901) and 0.5 mM ethylenediaminetetraacetic acid (EDTA) in phosphate-buffered saline (PBS) (1:1 mixture) at room temperature for 1 min. Then 2.3 × 10^5^ - 2.8 × 10^5^ cells were seeded on a Matrigel-coated 3.5 cm dish in the maintenance medium containing 5 µM ROCK inhibitor, Y27632 (Sigma, Y0503). The cells were maintained at 37 °C in a humidified atmosphere of 5% CO_2_. The medium was changed the next day into the maintenance medium without Y27632.

## Induction media

### From day 0 to day 2: PSM medium

The PSM medium is CDMi containing 10 μM CHIR99021 (Sigma, SML1046), 20 ng/ml bFGF, 10 μM SB431542 (Sigma, S4317), and 2 μM DMH1 (Sigma, D8946) as previously described^28,33,66^.

### From day 2 to day 6: Muscle medium_1

The basal medium is DMEM/F12 (Gibco, 11320033) containing 15% Knockout Serum Replacement (Gibco, 10828028), 0.1 mM nonessential amino acids (Gibco, 11140-035), 50 mM 2-mercaptoethanol (Gibco, 31350-010), and 0.5x penicillin/streptomycin (Gibco, 15140-122).

The muscle medium_1 is the basal medium supplemented with 10 ng/ml HGF (R&D, 2207-HG-025/CF), 2 ng/ml IGF (R&D, 791-MG-050), and 20 ng/ml bFGF.

### From day 2 to day 6: Muscle/Endothelial Cells (EC) medium_1

The muscle/EC medium_1 is the basal medium supplemented with 10 ng/ml HGF, 2 ng/ml IGF, 20 ng/ml bFGF, 2 µM forskolin (R&D, 2207-HG-025/CF), and 200 ng/ml VEGF165 (R&D, 2207-HG-025/CF).

### From day 6 to day 15: Muscle medium_2

The muscle medium_2 is the basal medium supplemented with 10 ng/ml HGF and 2 ng/ml IGF.

### From day 6 to day 15: Muscle/EC medium_2

The muscle/EC medium_2 is the basal medium supplemented with 10 ng/ml HGF, 2 ng/ml IGF, and 100 ng/ml VEGF165.

### Bovine muscle induction protocol in 2D

One day before the induction protocol started, 2.1 × 10^5^ - 2.5 × 10^5^ bovine ESCs were seeded on a Matrigel-coated 3.5 cm dish in the maintenance medium without IWR1 and with 5 μM Y27632. The maintenance medium was changed to the PSM medium the next day (defined as day 0). The cells were incubated in the PSM medium for 2 days, and the medium was changed every day. On day 2, the PSM medium was switched to the muscle medium_1, and the cells were incubated for 4 days with daily medium changes. On day 6, the muscle medium_1 was switched to the muscle medium_2, and the cells were incubated until day 15 with medium changes every other day.

### Bovine muscle and endothelial cell co-induction protocol in 2D

Bovine PSM cells were induced as described above. During days 2-6, the cells were incubated in the muscle/EC medium_1 with daily medium changes. During days 6-15, the cells were incubated in the muscle/EC medium_2 with medium changes every other day.

### Bovine muscle induction protocol in 3D

Bovine PSM cells were induced as described above. On day 2, PSM cells were trypsinized into single cells using TrypLE solution at 37 °C for 3 min. The cells were mechanically dissociated by pipetting and transferred into 1 ml of the muscle medium_1 containing 5 µM Y27632. The cell suspension was centrifuged at 258 xg for 3 min and resuspended into 1 ml of the muscle medium_1 containing Y27632. After one more wash with the muscle medium_1 containing Y27632, the supernatant was completely removed. The cell pellet was resuspended into 600-800 µl of the muscle medium_1 containing Y27632. The cell concentration was adjusted to 133.3 cells/µl with a sufficient amount of the pre-warmed muscle medium_1 containing Y27632. Then 150 µl of the cell suspension (= 20,000 cells) was aliquoted into each well of a U-Shaped-Bottom, 96-well-plate (96-well Clear Round Bottom Ultra-Low Attachment Microplate, Corning, 7007) by using a multichannel pipette. The 96-well plate was centrifuged at 258 xg for 2 min for the cells to settle down on the bottom of the plate. On day 3, 100 µl of the medium was carefully removed and replaced with 150 µl of the fresh muscle medium_1 containing 2% Matrigel (growth factor reduced, Corning, 356231). After day 4, the plate was placed on a rotary shaker (Celltron, INFORS HT) with the rotation speed set at 75 r.p.m. The medium was not changed until day 6. On day 6, the aggregates were collected using wide-bore tips and transferred into a 6-well plate containing 3 ml of the muscle medium_2. The aggregates were incubated in the muscle medium_2 until day 15 with medium changes every 3 days.

### Bovine muscle and endothelial cell co-induction protocol in 3D

The aggregate of 20,000 PSM cells was made as described above using the muscle/EC medium_1 instead of the muscle medium_1. During days 6-15, the aggregates were incubated in the muscle/EC medium_2 instead of the muscle medium_2.

### Immunohistochemistry (IHC) for 2D cell cultures

Cells in 2D cultures were washed twice with PBS(-) and fixed in 300 µl of 4% paraformaldehyde (PFA) at room temperature for 15 min. The cells were washed twice with PBSTw (0.2% Tween20 in PBS) for 5 min each, and then treated with the blocking buffer (PBS containing 3% Bovine Serum Albumin (BSA) and 0.2% Tween20) at room temperature for a few hours or at 4°C overnight. The cells were then incubated with the following primary antibodies in the blocking buffer at 4°C overnight: anti-TBX6 antibody (1/300 dilution, Abcam, ab38883), anti-SOX2 antibody (1/300 dilution, R&D, MAB2018), anti-BRACHYURY antibody (1/300 dilution, R&D, AF2085), anti-SARCOMERIC ALPHA ACTININ (SAA) antibody (1/1000 dilution, Abcam, ab9465), anti-BETA III TUBULIN (TUJ1) antibody (1/400 dilution, Abcam, ab18207), anti-MYOGENIN antibody (1/1000 dilution, Abcam, ab124800), anti-VE-CADHERIN antibody (1/300 dilution, Cell Signaling Technologies, 2500S), anti-Neurofilament heavy polypeptide (SMI-32) antibody (1/300 dilution, Abcam, ab8135), or anti-Myosin heavy chain (MHC) antibody (1/300 dilution, R&D, MAB4470). The next day, the cells were washed 3 times with PBSTw for 15 min each. Then the cells were incubated with the following secondary antibodies at a 1/500 dilution together with DAPI (1/500 dilution, Invitrogen, 62247) in PBSTw at 4°C overnight: Alexa Fluor 594 Goat anti-Rabbit IgG (H+L) (Invitrogen, A-11037), Alexa Fluor 647 Donkey anti-Goat IgG (H+L) (Invitrogen, A32849), or Alexa Fluor 488 Goat anti-Mouse IgG (H+L) (Invitrogen, A-11029). The cells were washed 3 times with PBSTw for 15 min each, and images were taken with an FV3000 confocal microscope (Olympus, FV3000 Fluoview RS software). Tiled images were stitched together using ImageJ Grid/Collection stitching. The fluorescent signal-positive area was measured using ImageJ thresholding.

### IHC for 3D aggregates

Aggregates in 3D cultures were collected using wide-bore tips into 2 ml Eppendorf tubes. After 2 washes with PBS(-), the aggregates were fixed in 500 μl of 4% PFA at 4°C overnight. They were washed 3 times with PBSTt (0.5% Triton X-100 in PBS) for 15 min each, and treated with the blocking buffer (PBS containing 5% Normal Goat Serum and 0.5% Triton X-100) at 4°C overnight on a shaker. Then the aggregates were incubated with the primary antibodies described above in the blocking buffer at 4°C for 2 days on a shaker. They were washed 3 times with PBSTt for 20 min each and incubated with the secondary antibodies described above in PBSTt at 4°C overnight on a shaker. After 3 washes with PBSTt for 20 minutes each, the aggregates underwent a clearing treatment. They were washed once with MilliQ and then embedded in 1% Agarose (low gelling temperature, Sigma, A9414) using a handmade rectangular box. The aggregates were then incubated with 100% Methanol in a glass bottle at room temperature for 2 days in the dark. The Methanol was changed daily. Then the Methanol was replaced with BABB (a mixture of Benzyl benzoate and Benzyl alcohol at a 1:1 ratio), and the aggregates were incubated at room temperature for 2 days in the dark. The BABB solution was changed daily. Images were taken using a MuVi-SPIM Light-Sheet Microscope with a clear chamber and BABB (Z = 2 μm, XY = 0.65 μm). Tiled images were stitched together using the MuVi-SPIM Light-Sheet Microscope’s software. The volume of the fluorescent signal-positive region was measured using the ilastik software with a pixel configuration and a custom-written Python script. Once the masked images of Z-stacks were generated by ilastik, the positive pixel values within each mask were counted, and these pixel counts were converted into a volume.

### Time-lapse imaging and quantification of HES7 oscillation

The HES7 promoter-luciferase reporter ESCs were incubated in the PSM medium containing 0.5 mM luciferin. The medium was changed daily. The luciferase activity was recorded using a Kronos Dio Luminometer (Atto, Kronos control software v2.3) every 10 min with 1 min exposure. The detrended signal was calculated by pyBOAT^67^.

### Calcium oscillation measurement

Calcium activity was visualized with Fluo-8 AM (Abcam, ab112129) following the manufacturer’s protocol. Briefly, cells were washed twice with PBS(-), and incubated with a mixture of Component A, B, and C at 37 °C for 30 min with or without 10 µM Tubocurarine Chloride Pentahydrate (Curare, Sigma, T2379). After 2 washes with the muscle medium_2, the Fluo-8 signals were recorded in the absence or presence of curare using the Thunder Imager Live Cell & 3D assay (Leica, LAS X software) every 100 msec.

For analyses, the Fluo-8 intensity from a randomly selected cell was measured using ImageJ. The signal was denoised using the Savitzky-Golay filter with a 50 window size using Python. The amplitudes between a signal peak and its adjacent two valleys were measured, and the peak was counted only if the smaller amplitude was larger than half of the larger amplitude. The total number of oscillation peaks in 2 min was manually counted.

### Quantitative RT-PCR (qRT-PCR)

Total RNA was extracted with RNeasy kit (Qiagen, 74104), and 1 µg RNA was reverse-transcribed with QuantiTect Reverse Transcription kit (Qiagen, 205311) to generate cDNA. The cDNA level was measured by a LightCycler 480 II (Roche) with LightCycler 480 SYBR Green I Master (Roche, 4707516001). The primer sequences are listed in Supplementary Table 1. The expression levels of the target genes were normalized by the *GAPDH* level.

### Endothelial network connectivity analysis

Endothelial network connectivity was analyzed with VascuMap software using a UNet architecture with a mixed vision transformer encoder^68^. The model, trained on endothelial cell images co-cultured with bovine muscle, labeled manually on an iPad 11 Pro with Photoshop and binarized via Python script. Images were divided into training and validation sets (70%/30%) with a fixed seed of 42. Training involved 200 epochs, batch size 16, RAdam optimizer, initial learning rate 0.001, step scheduler for rate reduction, and augmentations like flips and rotations. The loss function combined binary cross-entropy and dice coefficient, with the best IoU metric model on the validation set retained. For unseen data, VascuMap’s probability map was binarized using hysteresis thresholding (thresholds: 0.15, 0.5). The characterization of the binary masks returned the vessel area, total number of vessel segments, total number of branching points, and mean number of vessel segments per branch point.

## Acknowledgments

We are grateful to G. Martínez-Ara, J. Swoger, M. Coll Lladó, M. Costanzo, and M. Marchenko for imaging advice, to A. Akinbote and A. Olivares for discussion, to N. Gritti for analysis support, to the Mesoscopic Imaging Facility (MIF) at the European Molecular Biology Laboratory (EMBL), Olympus, and Luxendo/Bruker for imaging supports. Bovine ESCs were obtained from Dr. Juan Carlos Izpisua Belmonte^8^. This work was supported by EMBL; the Deutsche Forschungsgemeinschaft (DFG, German Research Foundation) under Germany’s Excellence Strategy - EXC 2068 - 390729961 - Cluster of Excellence Physics of Life of TU Dresden; the European Research Council (ERC) under the European Union’s Horizon 2020 research and innovation program (grant agreement No. 101002564 to M.E., 101040977 to K.H.). M.E. is supported by the Alexander von Humboldt Foundation in the framework of the Alexander von Humboldt Professorship endowed by the Federal Ministry of Education and Research. J.W. is a New York Stem Cell Foundation–Robertson Investigator and Virginia Murchison Linthicum Scholar in Medical Research. J.W. is supported by NYSCF, NIH (GM138565-01A1 and HD103627-01A1), Discovery and Innovation Grant from the American Society for Reproductive Medicine (ASRM) Research Institute, and The Welch Foundation (I-2088).

## Author contributions

M.E. conceived the work. M.S-M. performed most experiments and analyses. C.V. helped with experiments. L.R. and K.H. analyzed the endothelial networks. J.W. helped bovine ESC cultures. M.E. and M.S-M. wrote the manuscript, and all authors approved the final version.

---

**Supplementary Figure 1.**
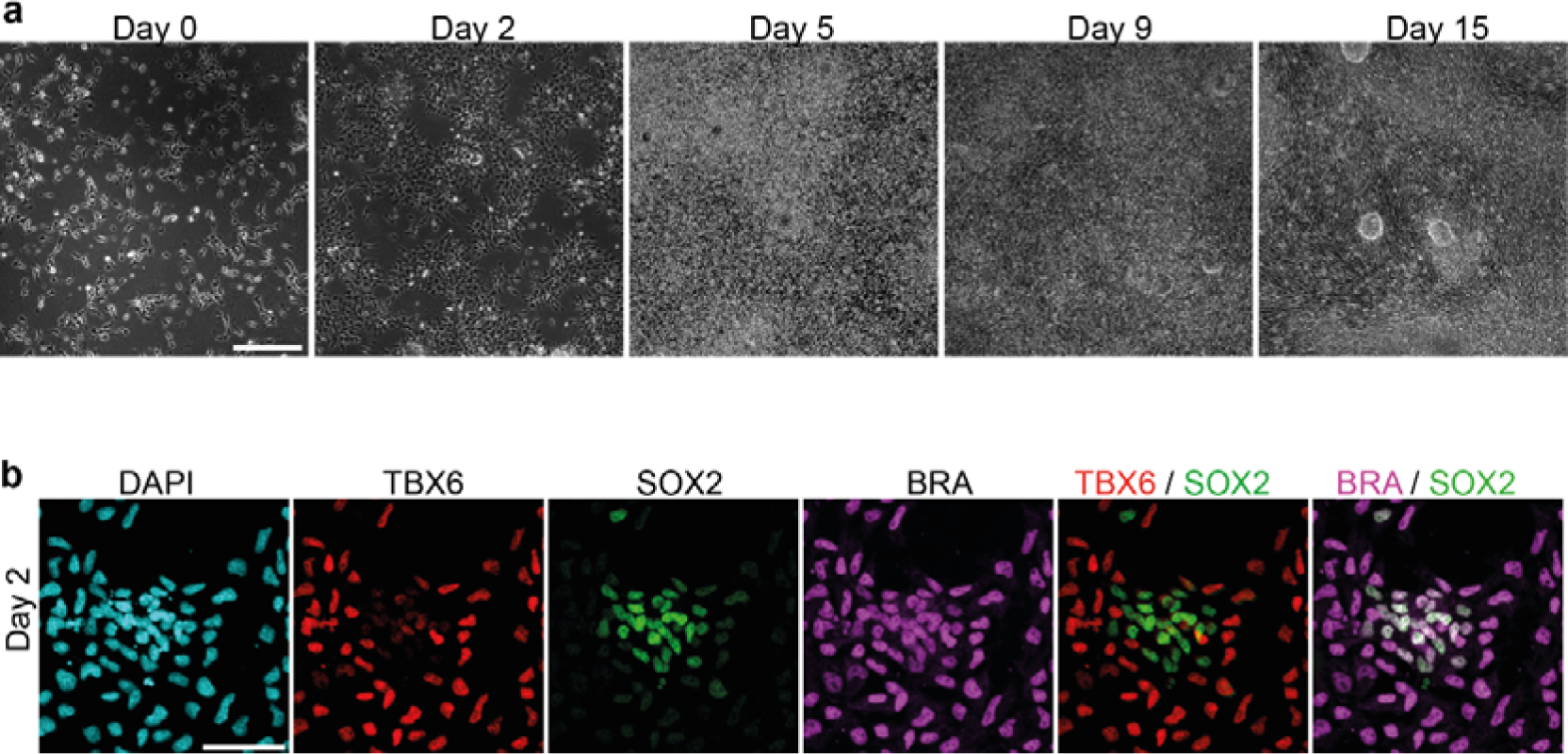
Induction of PSM and muscle fibers from bovine ESCs. **a,** Bright field images over the time course of skeletal muscle induction. The enlarged views of these images are shown in Fig. 1a. **b,** IHC images of the day-2 culture. The enlarged views of Fig. 1b. Scale bars: 300 µm (a) and 50 µm (b).

**Supplementary Figure 2.**
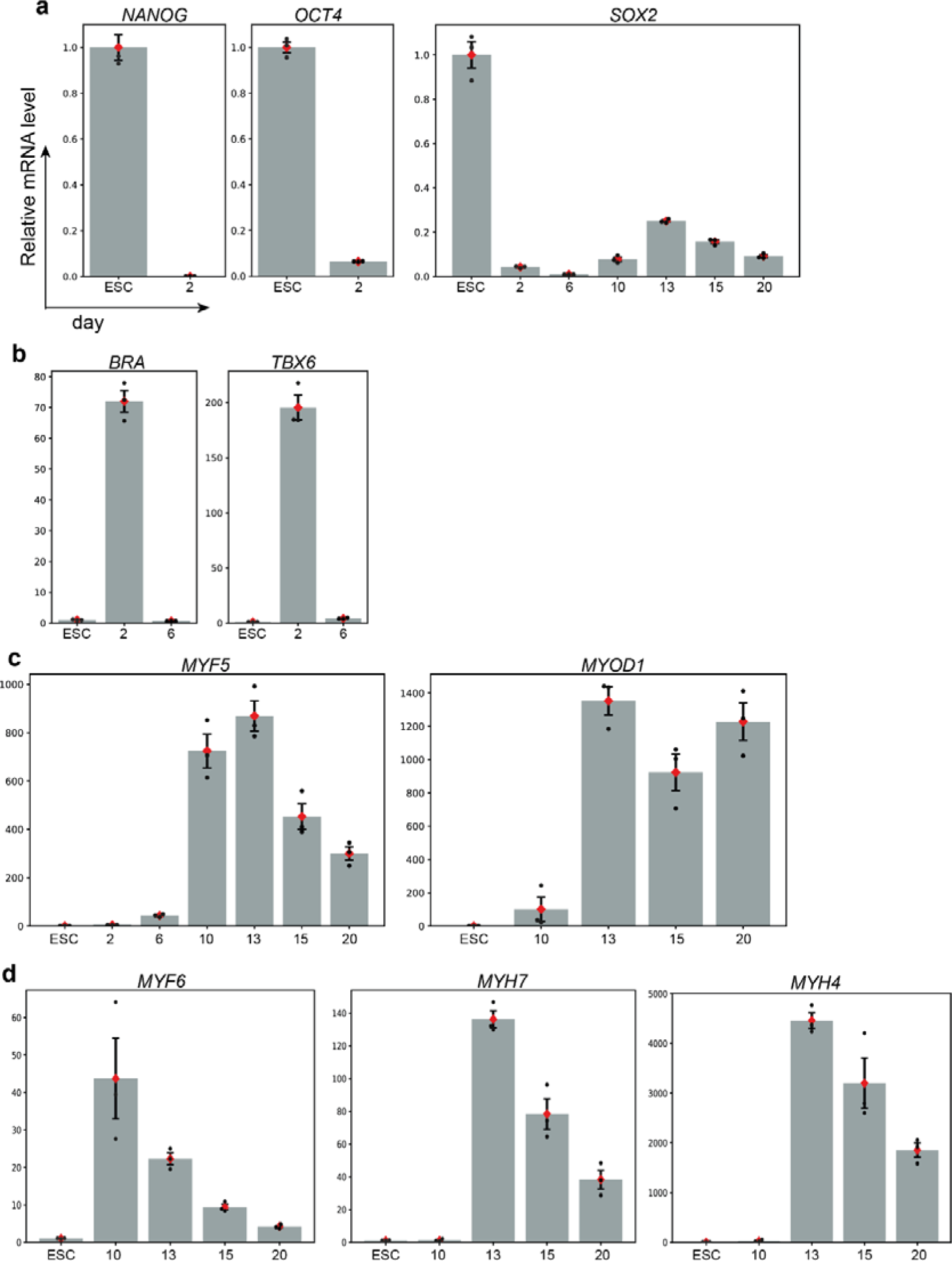
qRT-PCR measurements. Relative mRNA levels of selected marker genes in bovine ESCs and the time course muscle induction cultures. The values were normalized to *GAPDH* expression, and the average values of ESCs were set to 1. Mean ± SEM. N = 3 from 3 independent experiments. **a,** Pluripotency markers. **b,** PSM markers. **c,** Myoblast markers. **d,** Myocyte markers.

**Supplementary Figure 3.**
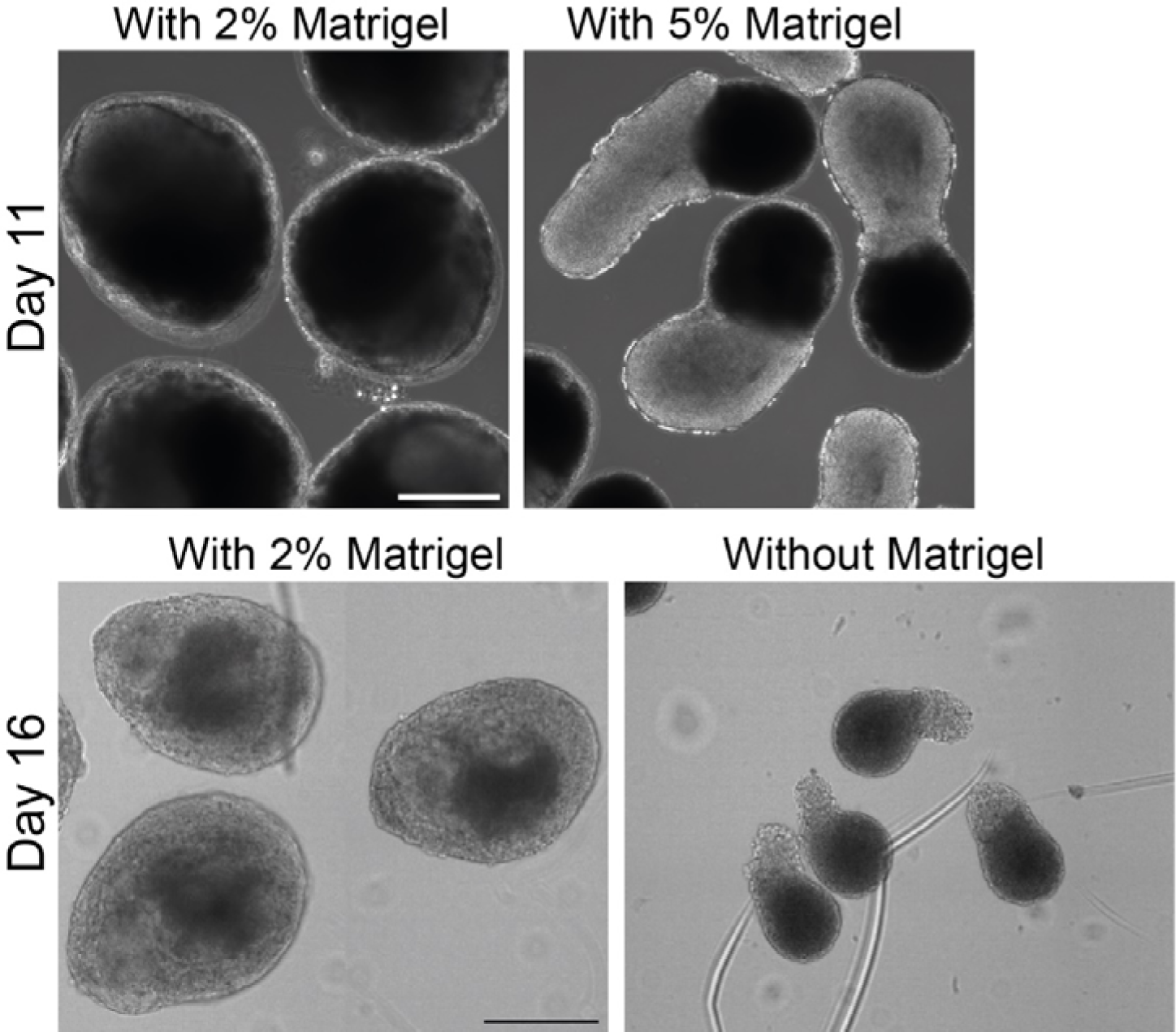
Effects of Matrigel on aggregates. Bright field images of the muscle aggregates made without or with (2% or 5%) Matrigel in the medium. 2% is the standard concentration. 3 independent experiments showed similar patterns. Scale bars: 300 µm.

**Supplementary Figure 4.**
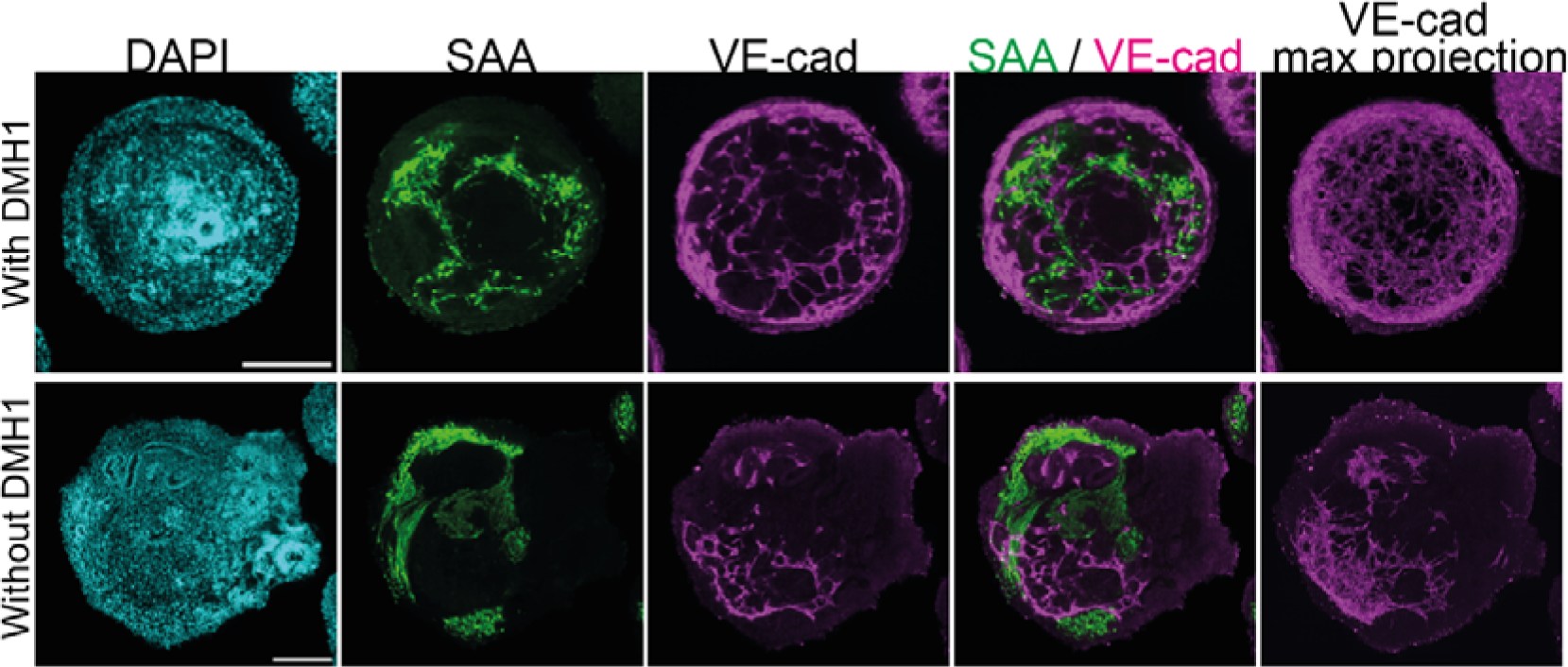
Effects of BMP inhibitor during PSM induction. IHC images of the day-15 aggregates. The aggregate made with the standard PSM medium with DMH1 (the BMP inhibitor) showed a uniform VE-cad-positive endothelial network throughout the tissue (top), while the one made without DMH1 showed local endothelial domains (bottom). The max projection of VE-cad used the Z-stacks within the 40 μm range around the central slice. 5-10 independent experiments showed similar patterns. Scale bars: 200 µm.

**Supplementary Table 1.**
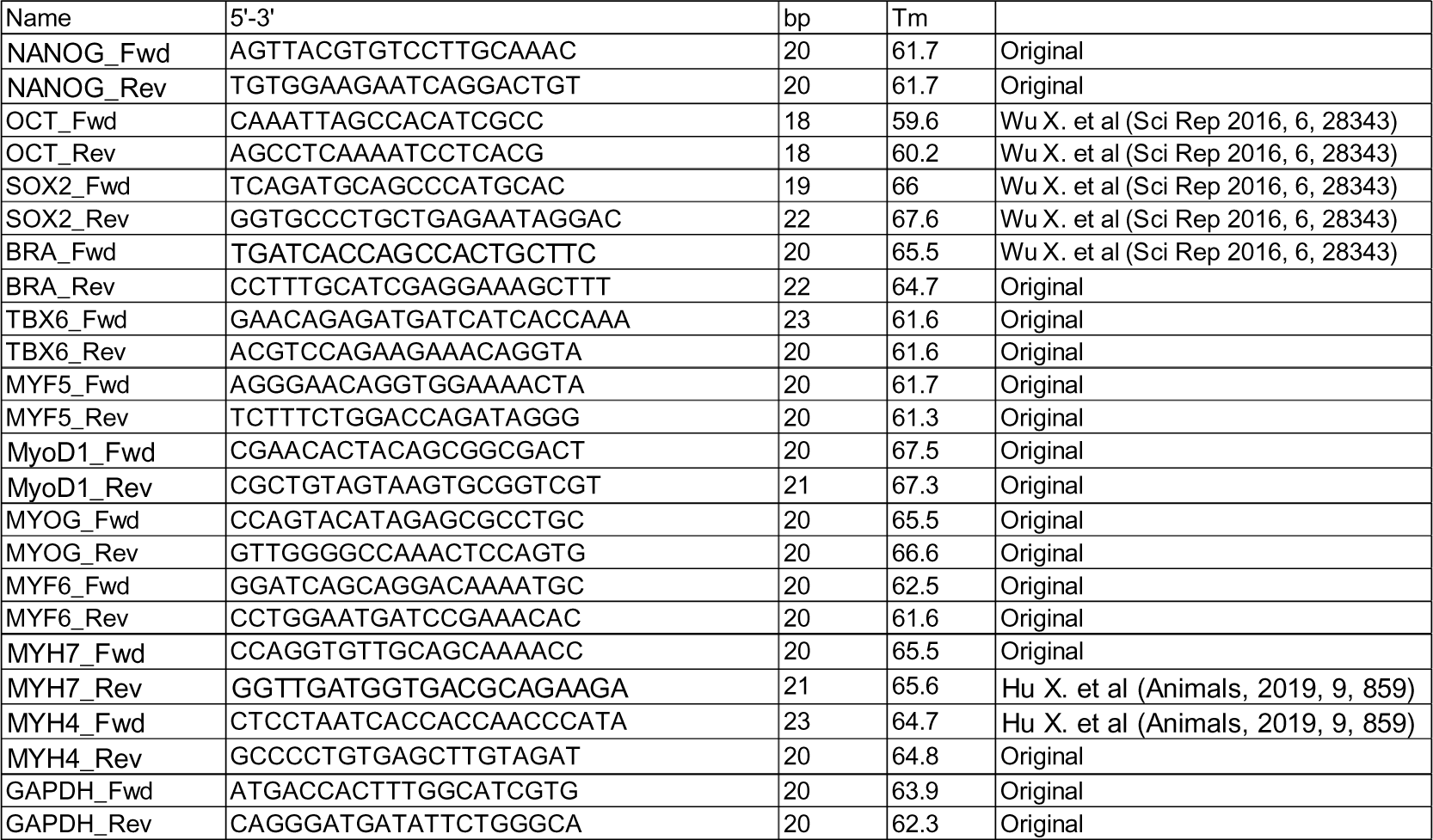
Primer sequences for qPCR.

**Supplementary Video 1**

Time-lapse imaging of calcium level fluctuations in the day-18 culture using Fluo-8 in the absence (left) and presence (right) of 10 μM curare. The time stamp indicates min:sec. The snapshot is also shown in Fig. 2a. Images were taken with the Thunder Imager Live Cell & 3D assay.

**Supplementary Video 2**

Z-stack IHC images of the day-15 aggregates from the 3D muscle and neuron co-induction protocol. SAA and TUJ1 are muscle fiber and neuronal markers, respectively. Z-stack interval is 2 μm. The snapshot is also shown in Fig. 4d. Images were taken by a MuVi-SPIM Light-Sheet Microscope.

**Supplementary Video 3**

Z-stack IHC images of the day-15 aggregate from the 3D muscle and endothelial co-induction protocol. SAA and VE-cad are muscle fiber and endothelial markers, respectively. Z-stack interval is 2 μm. The snapshot is also shown in Fig. 5d. Images were taken by a MuVi-SPIM Light-Sheet Microscope.

**Supplementary Video 4**

Z-stack IHC images of the day-15 aggregate from the 3D muscle and endothelial co-induction protocol. ZO-1 is a tight junction marker. Z-stack interval is 2 μm. The max projection is also shown in Fig. 5g. Images were taken by a MuVi-SPIM Light-Sheet Microscope.

**Supplementary Video 5**

Z-stack IHC images of the day-15 aggregate made from the PSM cells that were induced without BMP inhibitor DMH1. Z-stack interval is 2 μm. The snapshot is also shown in Supplementary Fig. 4. Images were taken by a MuVi-SPIM Light-Sheet Microscope.

